## Supplemental File 1 for "Systematic identification of molecular mediators underlying sensing of *Staphylococcus aureus* by *Pseudomonas aeruginosa*"

**TITLE**

Systematic identification of molecular mediators underlying sensing of *Staphylococcus aureus* by *Pseudomonas aeruginosa*

**AUTHORS**

Tiffany M. Zarrella[1, 2] and Anupama Khare[1, #]

[1]Laboratory of Molecular Biology, Center for Cancer Research, National Cancer Institute, National Institutes of Health, Bethesda, MD, USA

[2]Postdoctoral Research Associate Program, National Institute of General Medical Sciences, National Institutes of Health, Bethesda, MD, USA

### SUPPLEMENTAL FIGURES AND FIGURE LEGENDS

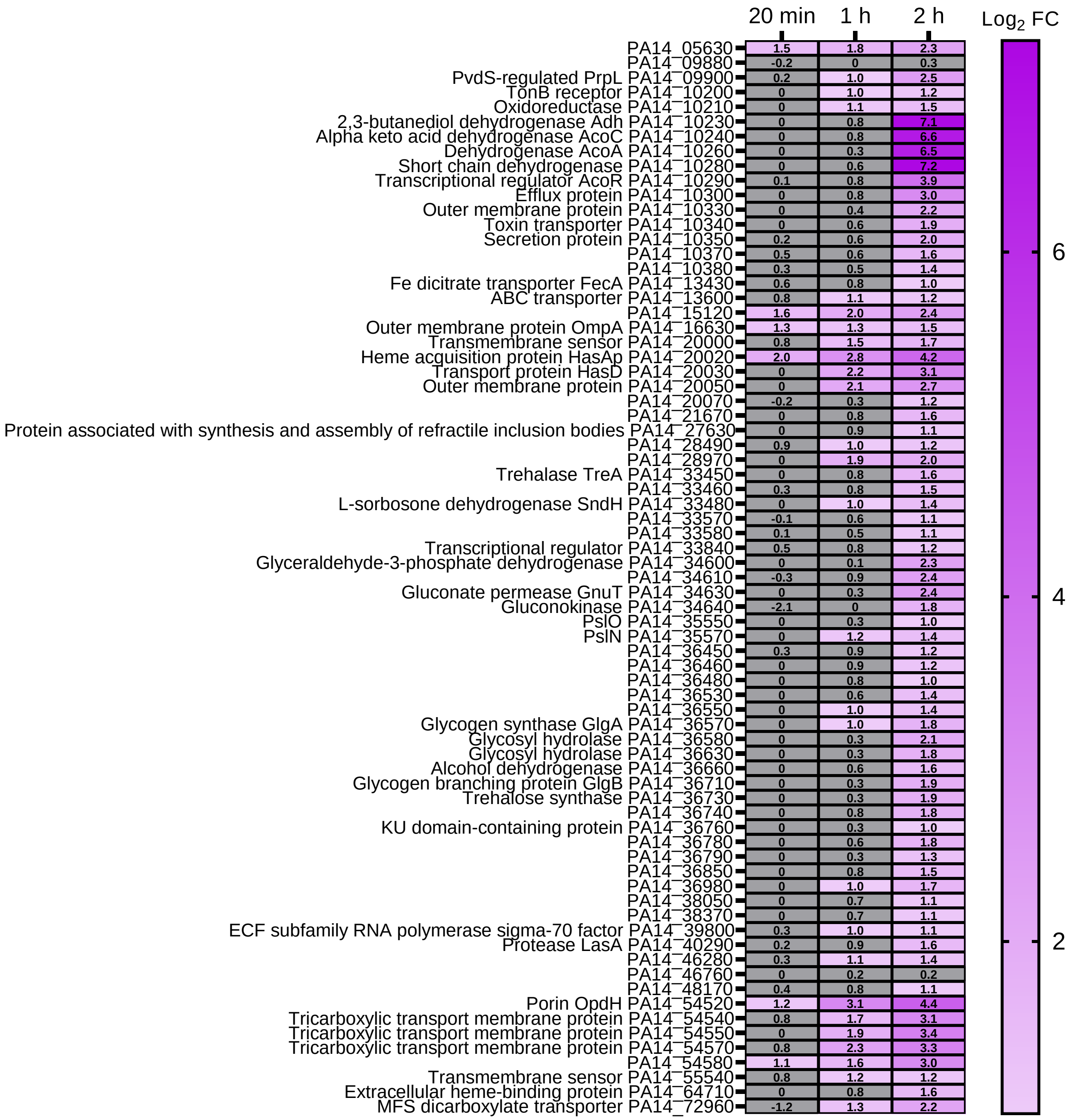


**FIGURE S1. Heat map of upregulated genes that increased in fold change over time upon addition of *S. aureus* supernatant.** The $\log_2$ fold changes for all upregulated genes with increasing fold change over time from 20 min to 1 h to 2 h are shown. Grey, $\log_2$ fold change < 1.

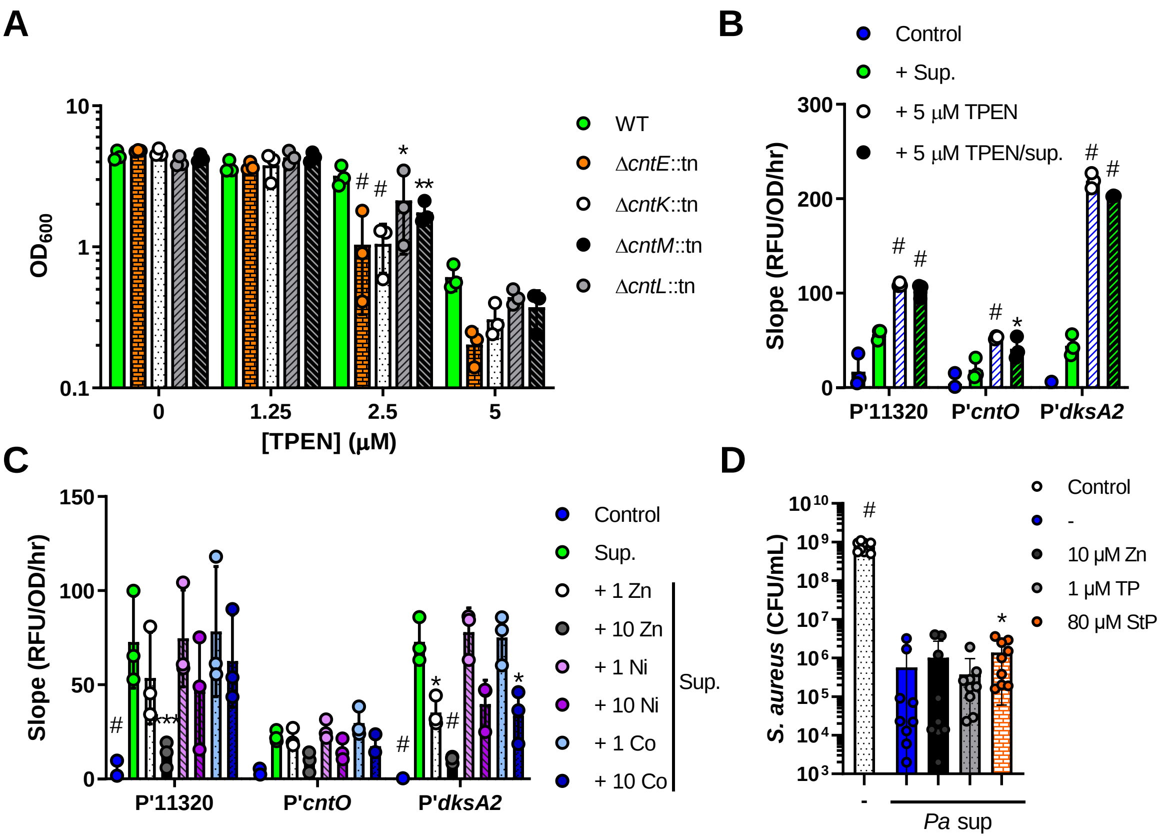


**FIGURE S2. Staphylopine and zinc levels affect *S. aureus* and *P. aeruginosa* physiology.** **(A)** WT *S. aureus* and NTML transposon mutants were grown in media with or without the addition of the zinc chelator TPEN at the concentrations indicated. Growth was measured at $OD_{600}$ after 24 h. Datasets were analyzed by a two-way ANOVA with Dunnett's test for multiple comparisons to the respective WT value. **(B-C)** RFU of mScarlet normalized to $OD_{600}$ over time after exposure to media control or *S. aureus* supernatant and/or addition of the indicated concentrations of **(B)** TPEN or **(C)** zinc, nickel, and cobalt in the indicated *P. aeruginosa* promoter-reporter strains. Datasets were analyzed by a two-way ANOVA with Tukey's test for multiple comparisons. Statistics shown represent tests comparing **(B)** the respective conditions/promoters with and without TPEN and **(C)** all treatments to the respective supernatant addition. **(D)** *S. aureus* cultures were grown in culture with 50% medium salts (Control) or cell-free supernatants of *P. aeruginosa* grown in the presence of the indicated additives. Colony forming units per milliliter of *S. aureus*

were calculated after 16 h of growth. Datasets were analyzed by one-way ANOVA with Dunnett's test for multiple comparisons to the *P. aeruginosa* supernatant sample. Data shown for all panels are the means of three independent biological replicates, including three separate supernatant collections in D. The error bars denote the SD. *, $p < 0.05$; **, $p < 0.01$; ***, $p < 0.001$; #, $p < 0.0001$.

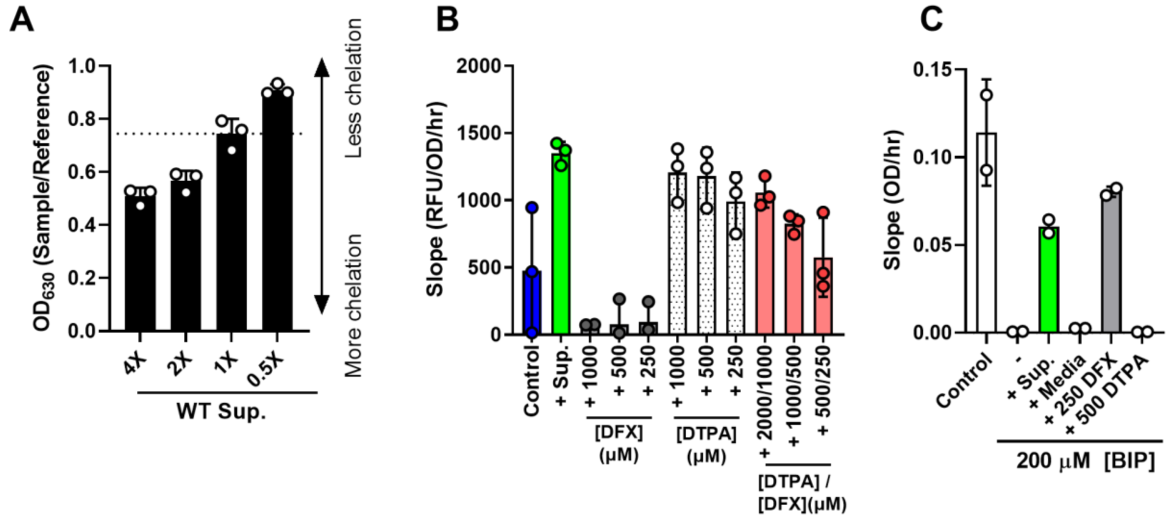


**FIGURE S3. Iron chelators affect induction of the *pvdG* promoter and iron delivery to *P. aeruginosa*.** **(A)** Chelation of different amounts of *S. aureus* supernatant was measured by CAS assay to construct a standard curve to calculate relative chelation. **(B)** RFU of mScarlet expressed from the promoter of *pvdG* normalized to $OD_{600}$ over time after exposure to media control, *S. aureus* supernatant, or the indicated concentrations of iron chelators DFX and DTPA. Data shown from three independent replicates. **(C)** Slope of $OD_{600}$ over time from 1-8 h of *P. aeruginosa* Δ*pvdJ* Δ*pchE* strain in medium control or medium with 200 µM BIP with the addition of whole WT *S. aureus* supernatant, media, or the iron chelators DFX or DTPA. Error bars denote the SD.

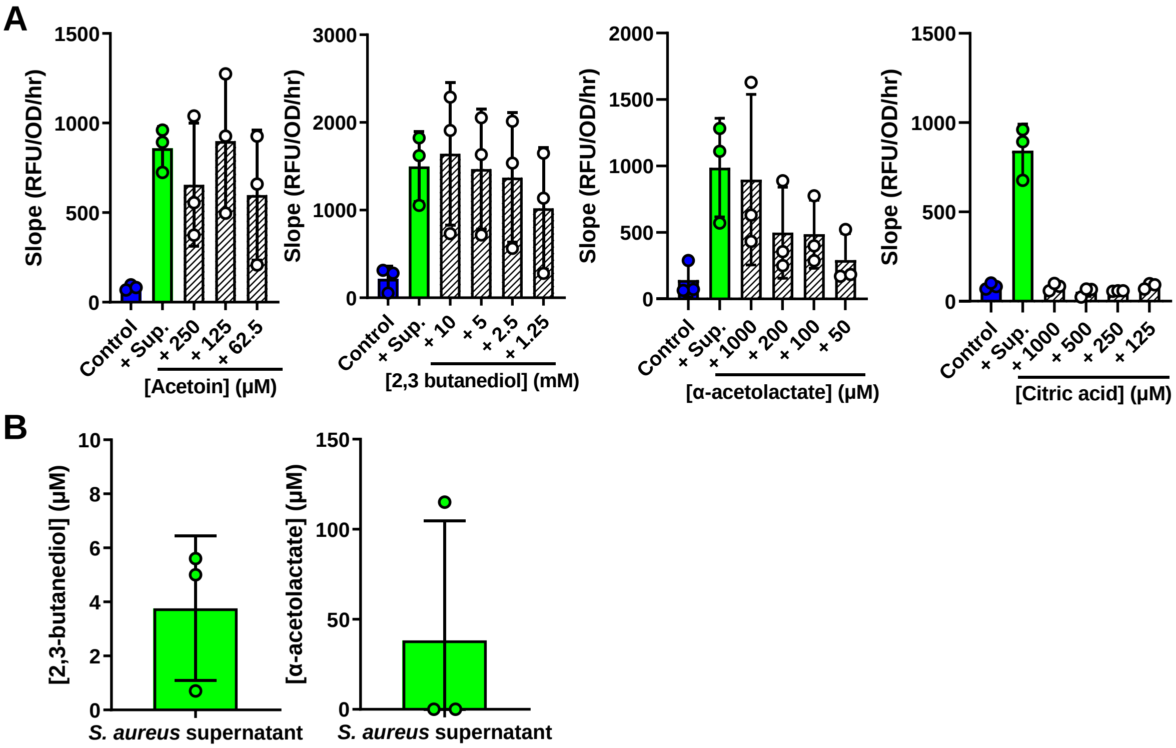


**FIGURE S4. The *acoR* promoter is induced by intermediate metabolites acetoin, 2,3-butanediol, and α-acetolactate but not citrate. (A)** RFU of mScarlet expressed from the *acoR* promoter normalized to $OD_{600}$ over time after exposure to media control, *S. aureus* supernatant, or addition of the indicated concentrations of each metabolite. **(B)** Quantification of 2,3-butanediol and α-acetolactate in *S. aureus* supernatant. Data shown for all panels is from three independent replicates. Error bars denote the SD.

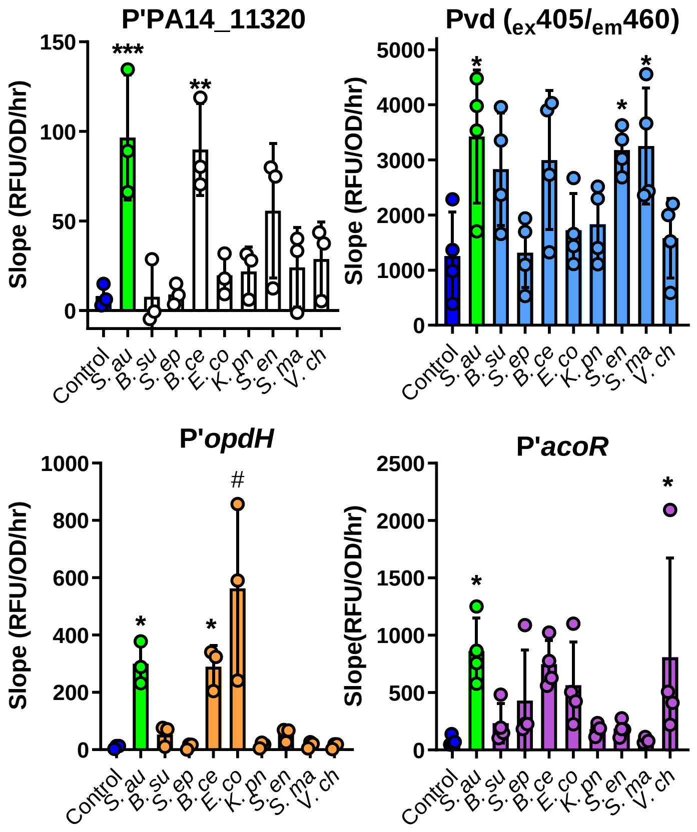


**FIGURE S5. Secreted products from diverse species induce response pathways in *P. aeruginosa*.** RFU of mScarlet expressed from the indicated promoter or pyoverdine (Pvd) normalized to $OD_{600}$ over time (promoter of PA14_11320, *opdH*, and *acoR* calculated from 1.5 to 5 h; Pvd calculated from 1 to 4 h) after exposure to media control, *S. aureus* supernatant, or supernatant from the indicated species: *Bacillus subtilis*, *Staphylococcus epidermidis*, *Burkholderia cenocepacia*, *Escherichia coli*, *Klebsiella pneumoniae*, *Salmonella enterica* Typhimurium, *Stenotrophomonas maltophilia*, and *Vibrio cholerae*. Data shown from at least three independent replicates. Error bars denote the SD. Datasets were analyzed by one-way ANOVA with Dunnett's test for multiple comparisons to the control. *, $p$ <0.05; **, $p$ <0.01; ***, $p$ <0.001; #, $p$ <0.0001.

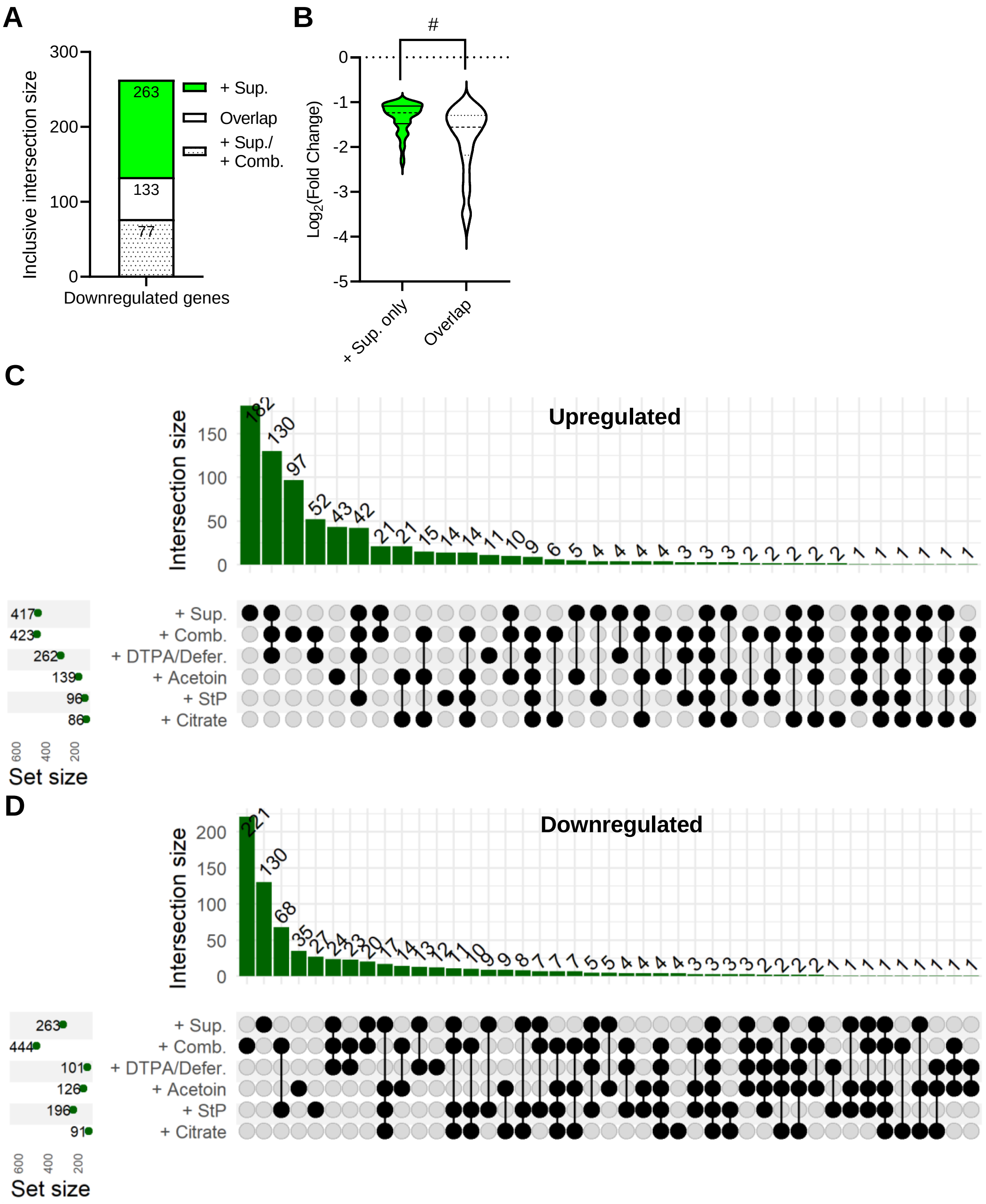


**FIGURE S6. Identified sensed products explain different proportions of the *P. aeruginosa* response to *S. aureus* supernatant. (A)** Inclusive intersection of all downregulated genes after addition of supernatant (+ Sup.) vs. at least one other condition (Overlap) or the combination of all molecules (+ Sup. / + Comb.). **(B)** $Log_2$ fold change of the downregulated genes that are non-intersecting (+ Sup. only) or intersecting among supernatant and at least one other condition (Overlap). Datasets were analyzed by a two-tailed *t*-test. #, $p < 0.0001$. **(C and D)** UpSet plot showing all exclusive intersections of **(C)** upregulated or **(D)** downregulated genes at 20 min and 2 h after addition of *S. aureus* supernatant, the indicated products, or the combination of all products (+ Comb.). All intersections are shown.

### SUPPLEMENTAL TABLES

**Table S1. Gene expression in *P. aeruginosa* at different time points after exposure to *S. aureus* supernatant or media control determined from RNA-seq analysis. (Excel)** Gene expression was analyzed by RNA-seq from RNA purified from two biological replicates of each treatment at each time point (20 min, 1 h, and 2 h after addition of *S. aureus* supernatant or media control).

**Table S2. Differentially expressed genes between *P. aeruginosa* after exposure to *S. aureus* supernatant or media control determined from RNA-seq analysis. (Excel)** Gene expression was analyzed by RNA-seq from RNA purified from two biological replicates of each treatment at each time point (20 min, 1 h, and 2 h after addition of *S. aureus* supernatant or media control). $Log_2$ fold change compared to the media control is shown.

**Table S3. Gene ontology enrichment of upregulated and downregulated genes after *P. aeruginosa* exposure to *S. aureus* exoproducts. (Excel)** PANTHER Overrepresentation Test (Released 20210224). GO Ontology database DOI: 10.5281/zenodo.5228828 Released 2021-08-18. Fisher's exact test with Bonferroni correction.

**Table S4. Differential expression of genes after 20 min in the Fur, PvdS, and Zur regulons. (Excel)** Fur and Iron Starvation (IS) box genes as listed in Ochsner et al. (1). Zur box genes as listed in Pederick et al. (2). $Log_2$ fold change compared to the media control is shown.

**Table S5. Slopes of promoter induction from NTML screen with the P'PA14_11320 promoter-reporter strain. (Excel)**

**Table S6. Slopes of promoter induction from NTML screen with the P'*pvdG* promoter-reporter strain. (Excel)**

**Table S7. Iron chelating activity of upregulating and downregulating supernatants. (Excel)** Chelating activity was determined by CAS assay and normalized to WT supernatant (relative chelation).

**Table S8. Slopes of promoter induction from NTML screen with the P'*opdH* promoter-reporter strain. (Excel)**

**Table S9. Gene ontology enrichment of NTML mutants that upregulate or downregulate P'*opdH* in *P. aeruginosa*. (Excel)** PANTHER Overrepresentation Test (Released 20200728). GO Ontology database DOI: 10.5281/zenodo.3980761 Released 2020-08-10. Fisher's exact test with Bonferroni correction.

**Table S10. Citrate measurements in upregulating and downregulating supernatants. (Excel)**

**Table S11. Slopes of promoter induction from NTML screen with the P'*acoR* promoter-reporter strain. (Excel)**

**Table S12. Acetoin measurements in upregulating and downregulating supernatants. (Excel)**

**Table S13. Differential expression of genes in *P. aeruginosa* after exposure to *S. aureus* supernatant or the identified products. (Excel)** Gene expression was analyzed by RNA-seq from RNA purified from two biological replicates of each treatment at each time point of 20 min and 2 h. $Log_2$ fold change was calculated compared to the media control.

**Table S14. Bacterial strains and plasmids used in this study.**

| Name | Description[a] | Source |
|---|---|---|
| **Strains** | | |
| *P. aeruginosa* | | |
| PA14 | University of California Berkeley Plant Pathology (UCBPP)-PA14 | (3) |
| AK625 | Δ*pvdJ* Δ*pchE* | (4) |
| SB89 | PA14 P'*dksA2*-mScarlet (pSB83) unmarked | This study |
| SB91 | PA14 P'11320-mScarlet (pSB85) unmarked | This study |
| SB124 | PA14 P'*opdH*-mScarlet (pSB109) unmarked | This study |
| SB136 | PA14 P'*acoR*-mScarlet (pSB121) unmarked | This study |
| SB141 | PA14 Δ*cntO* P'11320-mScarlet (pSB85) unmarked | This study |
| SB142 | PA14 Δ*cntI* P'11320-mScarlet (pSB85) unmarked | This study |
| SB143 | PA14 Δ*cnt* P'11320-mScarlet (pSB85) unmarked | This study |
| SB204 | PA14 P'*pvdG*-mScarlet (pSB175) unmarked | This study |
| *S. aureus* | | |
| JE2 | *Staphylococcus aureus* subsp. *aureus* USA300_FPR3757 (CA-MRSA)-JE2 | (5) |
| CF049 | AMT0150-13 | CFF Isolate Core |
| CF061 | AMT0150-28 | CFF Isolate Core |
| CF085 | AMT0458-3; co-isolated with *P. aeruginosa* | CFF Isolate Core |
| CF089 | AMT0461-13; co-isolated with *P. aeruginosa* | CFF Isolate Core |
| *E. coli* | | |
| *ccdB* Survival 2 $T1^R$ | *E. coli* strain used for maintenance of pDONR plasmid | Invitrogen |
| DH5ɑ | *E. coli* strain used for cloning | NEB |
| AK111 | MG1655 SB144 | (6) |
| S17-1 λ-pir | *E. coli* strain used for conjugation | (7) |
| Other species | | |
| SB80 | *Staphylococcus epidermidis* (Winslow and Winslow) Evans FDA strain PCI 1200; ATCC 12228 | ATCC |
| SB81 | *Salmonella enterica* subsp. *Enterica* (ex Kauffmann and Edwards) Le Minor and Popoff serovar Typhimurium; ATCC 29630 | ATCC |
| SB145 | *Bacillus subtilis* PY79 | K. Ramamurthi |
| SB146 | *Burkholderia cenocepacia*; ATCC 25608 | S. Adhya |
| SB147 | *Klebsiella pneumoniae* subsp. *pneumoniae* KPNIH1 | S. Adhya |
| SB148 | *Stenotrophomonas maltophilia* (Hugh) Palleroni and Bradbury K279a; ATCC BAA-2423 | ATCC |
| SB149 | *Vibrio cholerae* | S. Adhya |
| **Plasmids** | | |
| *Promoter-reporter* | | |
| pSEK109 | pLD3208. Shuttle vector with FRT sites and *mScarlet* ORF; $Tet^r$ $Gent^r$ | (8) |
| pSB83 | pSEK109: P'PA14_73020 (*dksA2*) | This study |
| pSB85 | pSEK109: P'PA14_11320 | This study |
| pSB109 | pSEK109: P'PA14_54520 (*opdH*) | This study |

| | | |
|---|---|---|
| pSB118 | pSEK109: P'PA14_63960 (*cnt*) | This study |
| pSB121 | pSEK109: P'PA14_10290 (*acoR*) | This study |
| pSB175 | pSEK109: P'PA14_33270 (*pvdG*) | This study |
| *Gene deletion* | | |
| pDONRP EX18Gm | Shuttle vector with *attP* sites and *ccdB*; Cm$^r$ Gent$^r$ | (9) |
| pSB138 | pDONRPEX18Gm: ΔPA14_63960 (*cntO*) | This study |
| pSB139 | pDONRPEX18Gm: ΔPA14_63910 (*cntI*) | This study |
| pSB140 | pDONRPEX18Gm: ΔPA14_63960-PA14_63910 (*cnt*) | This study |
| *Remove antibiotic resistance cassette* | | |
| pFLP2 | Expressing Flp recombinase; Ap$^r$ Car$^r$ | (10) |

[a] Ap$^r$, ampicillin resistance (*E. coli*); Car$^r$, carbenicillin resistance (*P. aeruginosa*); Cm$^r$, chloramphenicol resistance (*E. coli*); Gent$^r$, gentamicin resistance (*P. aeruginosa*); Tet$^r$, tetracycline resistance (*E. coli*)

**Table S15. Primers used in this study.**

| Primer | Sequence 5′→3′ | Site^ | Location* | Application |
|---|---|---|---|---|
| Pa020 | atcccgacgggcccggtaccactagtcaggtaagcgcattccgc | SpeI | F P'PA14_73020 | Promoter-reporter |
| Pa024 | atcccgacgggcccggtaccactagtcagctggcgccaatcctc | SpeI | F P'PA14_11320 | Promoter-reporter |
| Pa029 | atcccgacgggcccggtaccactagtcgggaagacagggagaaatc | SpeI | F P'PA14_10290 | Promoter-reporter |
| Pa031 | atcccgacgggcccggtaccactagtgcgagctgcggatgatct | SpeI | F P'PA14_54520 | Promoter-reporter |
| Pa045 | atcccgacgggcccggtaccactagtgcgggaggaattggagat | SpeI | F P'PA14_63960 | Promoter-reporter |
| Pa094[#] | ccgttcatagaaccctccatg | - | R mScarlet | Sanger sequencing |
| Pa110 | atctccttcttaaatctagactcgaggggctgaacctgggaaat | XhoI | R P'PA14_73020 | Promoter-reporter |
| Pa112 | atctccttcttaaatctagactcgaggaacgggactccggcaag | XhoI | R P'PA14_11320 | Promoter-reporter |
| Pa113 | atctccttcttaaatctagactcgaggtccacatggtccttcgagt | XhoI | R P'PA14_10290 | Promoter-reporter |
| Pa114 | atctccttcttaaatctagactcgagggaagtcgacatgcagtggt | XhoI | R P'PA14_54520 | Promoter-reporter |
| Pa121 | atctccttcttaaatctagactcgagaccgaggacaagcgacac | XhoI | R P'PA14_63960 | Promoter-reporter |
| Pa127 | ggggacaagtttgtacaaaaaagcaggctcacgaagttcaccagggtcagt | *attB1* | Up PA14_63960 | Generate mutant |
| Pa128 | ggcggtagagggctcagtacatgggaaatcgcaccag | - | Up and down PA14_63960 | Generate mutant |
| Pa129 | ctggtgcgatttcccatgtactgagccctctaccgcc | - | Up and down PA14_63960 | Generate mutant |
| Pa130 | ggggaccactttgtacaagaaagctgggtaacaaatgtgcgccgacct | *attB2* | Down PA14_63960 | Generate mutant |
| Pa131[#] | gaaatgcagcggatcgag | - | Up PA14_63960 | Sanger sequencing |
| Pa132[#] | agtaccgcgcttttctgct | - | Down PA14_63960 | Sanger sequencing |
| Pa133 | ggggacaagtttgtacaaaaaagcaggctcagtgcgagctgagcgaact | *attB1* | Up PA14_63910 | Generate mutant |
| Pa134 | ggaggctcagcccttcttcttcagcaggtcgagcac | - | Up and down PA14_63910 | Generate mutant |
| Pa135 | gtgctcgacctgctgaagaagaagggctgagcctcc | - | Up and down PA14_63910 | Generate mutant |
| Pa136 | ggggaccactttgtacaagaaagctgggtagctgctctacagcatctcgac | *attB2* | Down PA14_63910 | Generate mutant |
| Pa137[#] | ccggatactttccgagcag | - | Up PA14_63910 | Sanger sequencing |
| Pa138[#] | acggcagtccatggaagat | - | Down PA14_63910 | Sanger sequencing |

| | | | | |
|---|---|---|---|---|
| Pa139 | ggaggctcagcccttcttcatgggaaatcgcaccag | - | Down PA14_63910 and up PA14_63960 | Generate mutant |
| Pa140 | ctggtgcgatttcccatgaagaagggctgagcctcc | - | Up PA14_63960 and down PA14_63910 | Generate mutant |
| Pa157 | atcccgacgggcccggtaccactagtggaaatcaccttgctgcgg | SpeI | F P'PA14_33270 | Promoter-reporter |
| Pa158 | atctccttcttaaatctagactcgagctgccgacctccttgcg | XhoI | R P'PA14_33270 | Promoter-reporter |

* F = forward of ORF; R = reverse of ORF; up = upstream arm of gene; down = downstream arm of gene.

[#] Primers utilized for Sanger sequencing.

^ Site is underlined in primer sequence.
